# Specific inhibition of α-synuclein oligomer generation and toxicity by the chaperone domain Bri2 BRICHOS

**DOI:** 10.1101/2023.12.07.570346

**Authors:** Laurène Adam, Rakesh Kumar, Luis Enrique Arroyo-Garcia, Willem Hendrik Molenkamp, Jan Stanislaw Nowak, Hannah Klute, Azad Farzadfard, Rami Alkenayeh, Janni Nielsen, Henrik Biverstål, Daniel E. Otzen, Jan Johansson, Axel Abelein

## Abstract

Understanding the molecular mechanisms of neurodegenerative diseases and finding efficient treatments have been major priorities for research and society, yet new therapeutic approaches remain essential to face the socio-economic burden caused by these devastating diseases. Protein misfolding and aggregation are involved in several neurodegenerative disorders, such as α-synuclein (αSyn) implicated in Parkinson’s disease. Elucidating the microscopic nucleation mechanisms has opened new opportunities to develop therapeutics against toxic mechanisms and species. Here, we show that naturally occurring molecular chaperones, represented by the anti-amyloid Bri2 BRICHOS domain, can be used to target αSyn-associated nucleation processes and structural species related to neurotoxicity. Our findings revealed that BRICHOS predominately suppresses the formation of new nucleation units on the fibrils surface (secondary nucleation), in addition to fibril-end elongation. This mechanism implies a drastic decrease of the oligomer generation rate. Besides targeting secondary nucleation sites on the fibril surface, BRICHOS directly binds to oligomeric αSyn species. Further, using *ex vivo* experiments, BRICHOS effectively diminishes αSyn fibril-related toxicity to hippocampal electrophysiology. Our studies show that molecular chaperones can be utilized as tools to target molecular processes and structural species related to αSyn neurotoxicity and have the potential as protein-based treatments against neurodegenerative disorders.

## INTRODUCTION

Protein misfolding and self-assembly have been linked to several neurodegenerative disorders where the formation of amyloid fibrils is closely associated to the pathologies of the disease pathology (Knowles et al., 2014). Amyloid formation of the α-synuclein protein (αSyn) is firmly linked to several neurodegenerative diseases, called α-synucleinopathies, including Parkinson’s disease (PD), dementia with Lewy bodies and multiple system atrophy. These disorders are characterized by abnormal accumulation of misfolded αSyn into intracellular inclusions in neurons called Lewy bodies (Spillantini et al., 1998). Mutations in the gene coding for αSyn cause a different aggregation mechanism, therefore an early onset and/or more aggressive forms of PD (such as A30P, A53T, E46K, H50Q and G51D), providing strong evidence that αSyn aggregation plays a crucial role in PD (Kara et al., 2013). Immense efforts have been undertaken to suppress amyloid formation but no effective treatments against α-synucleinopathies have been developed so far. In particular, strategies to inhibit toxic aggregation pathways and species are demanded, since the generation of intermediate, oligomeric or low molecular weight fibrillar αSyn assemblies have been suggested to be the cause of αSyn-associated neurotoxic effects (Winner et al., 2011).

Molecular chaperones are nature’s own way to counteract protein misfolding. Several chaperone families have been identified, that in addition to the classical chaperone functions target amyloids (Abelein & Johansson, 2023; Wentink et al., 2019). A good example is provided by the endoplasmic reticulum (ER) luminal chaperone domain BRICHOS from the transmembrane Bri2 protein in which mutations cause familial British and Danish dementia (Leppert et al., 2023). The recombinant human (rh) Bri2 BRICHOS protein delays amyloid-β (Aβ) aggregation, the aggregation-prone peptide involved in Alzheimer’s disease (Arosio et al., 2016; Chen et al., 2017; Chen et al., 2020; Zhong et al., 2022). Monomeric Bri2 BRICHOS and specifically the single-point mutant R221E Bri2 BRICHOS (referred to as BRICHOS below), which forms stable monomers, were shown to suppress Aβ42 fibrillization and reduce Aβ42-related toxicity to mouse hippocampal slices in electrophysiological measurements (Chen et al., 2017; Chen et al., 2020). This inhibitory mechanism on Aβ42 fibrillization is likely mediated through the binding of BRICHOS to the surface of Aβ42 fibrils, which prevents the formation of potentially neurotoxic oligomeric or low molecular weight fibrillar species (Chen et al., 2017; Chen et al., 2020). Notably, BRICHOS exhibited crucial abilities of a potential drug candidate by crossing the blood-brain barrier in mice (Galan-Acosta et al., 2020; Manchanda et al., 2022; Tambaro et al., 2019). Further, amyloid precursor protein (APP) knock-in mouse models treated with rh R221E Bri2 BRICHOS monomers exhibited reduced Aβ plaque content and improved cognitive and memory function compared to controls (Manchanda et al., 2022). Hence, based on these studies, Bri2 BRICHOS showed efficient treatment of AD pathology in relevant *in vivo* models.

Of note, besides Aβ, BRICHOS (fused to mCherry) can bind several other types of amyloid fibrils like islet amyloid polypeptide (IAPP), the designed β-structure protein β17 and αSyn. This points toward a generic ability of BRICHOS to bind to amyloid surfaces (Biverstål et al., 2020). The fibril surface has been identified as a catalyzer for secondary nucleation processes, *i.e.* the formation of new nucleation units on the fibril surface. The auto-catalytic process of secondary nucleation was found to be the dominant mechanism in Aβ and αSyn self-assembly (Buell et al., 2014; Cohen et al., 2013; Meisl et al., 2014). This process is also the major source of the generation of new oligomeric or small fibrillar species, and hence potentially closely linked to neurotoxic effects (Abelein & Johansson, 2023; Cohen et al., 2015). Specifically targeting secondary nucleation and the species it generates may hence enable efficient therapeutic approaches.

Here, we report that BRICHOS efficiently delays αSyn self-assembly by binding to the fibril surface and thereby particularly suppresses secondary nucleation reactions, in addition to fibril-end elongation. This inhibitory mechanism drastically reduces the rate of formation of new nucleation units, which might otherwise convert to potentially neurotoxic oligomeric and/or small fibrillar assemblies. Furthermore, our findings give evidence that BRICHOS directly binds to oligomeric species and prevents toxic effects associated to αSyn. Hence, the current study reveals molecular insights into how αSyn aggregation and toxicity can be targeted by a molecular chaperone domain. This opens possibilities for developing new treatment approaches against amyloid disease.

## RESULTS

### BRICHOS efficiently retards αSyn bulk aggregation

We first followed the fibrillization kinetics of 70 μM wildtype (WT) αSyn applying a Thioflavin T (ThT) assay (using agitation with glass beads at physiological pH 7.4) in the presence of different concentration of recombinant human (rh) Bri2 BRICHOS R221E monomers. We observed that the aggregation of αSyn was delayed upon addition of BRICHOS in a concentration-dependent manner, resulting in a delay of the aggregation half-times from 18.3 ± 1.8 h to 53.1 ± 3.0 h (at 95 % molar equivalent of BRICHOS), obtained by sigmoidal fits to the individual aggregation traces (Fig. 1a). At higher BRICHOS concentrations, the inhibitory effect appears to be saturated, and no further retardation of the aggregation is observed in the range of 25 % to 100 % molar equivalent of BRICHOS (Fig. 1b). A similar phenomenon was previously reported for the Aβ42 peptide, where a saturation of bound BRICHOS on to the fibril surface was suggested (Cohen et al., 2015).

**Figure 1:**
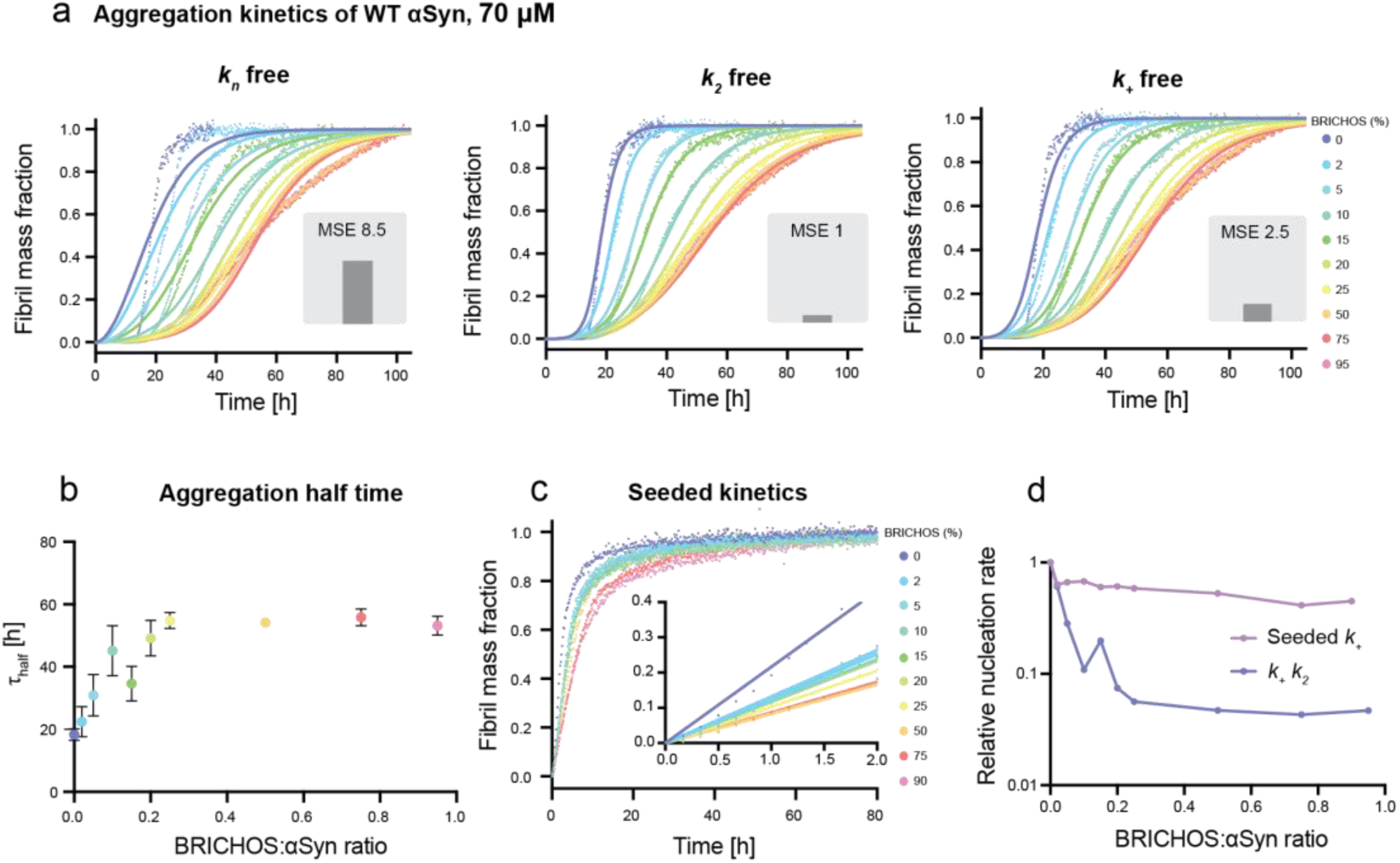
BRICHOS inhibits WT αSyn aggregation by reducing predominately secondary nucleation in addition to fibril-end elongation. **(a)** ThT fluorescence aggregation kinetics of 70 µM WT αSyn in 20 mM NaP buffer, 0.2 mM EDTA at pH 7.4, using agitation and glass beads, in the presence of 0 to 95 % BRICHOS (color gradient from blue to red) at 37°C fitted to a secondary nucleation dominated kinetic model with AmyloFit (Meisl et al., 2016). Individual fits (solid lines) to normalized and averaged aggregation traces (dots) of six replicates of αSyn aggregation kinetics are shown in the presence of BRICHOS where either *k_n_*, *k_2_* or *k_+_* is the free fitting parameter. The normalized mean square error (MSE) value refers to the global fits, where the *k_2_* is the free fitting parameter that best describes the data followed by *k_+_* and *k_n_*. **(b)** Aggregation half-time (τ_half_), revealing an increased value with increasing BRICHOS:αSyn ratio, which reaches a plateau at higher molecular ratios. The errors reflect the standard derivation of six replicates. **(c)** Highly seeded αSyn aggregation kinetics using 5 % seeds of sonicated αSyn fibrils and 0 to 90 % BRICHOS (color gradient from blue to red). **(d)** Combined rate constants *k_+_k_2_* (where *k_+_k_n_* is fixed as a global fit constant) and *k_+_* extracted from highly seeded kinetics plotted against the BRICHOS:αSyn ratio, showing a predominate inhibitory effect of BRICHOS on *k_2_* compared to *k_+_*.

To test whether the inhibitory effect originates from electrostatic interactions between BRICHOS and αSyn, we further conducted the aggregation kinetics in the presence of 154 mM NaCl, *i.e.* close to physiological salt concentration (SI Fig. S1). We observed that, in the presence of salt, the aggregation kinetics of αSyn is greatly accelerated and the aggregation half time is reduced to 9.9 ± 0.7 h. This effect with accelerated kinetics in the presence of salt has previously been reported (Fujiwara et al., 2019; Havemeister et al., 2023; Munishkina et al., 2004). Notably, also under these conditions BRICHOS efficiently retards the aggregation process, delaying the aggregation half time to 36.5 ± 6.6 h at 90 % molar equivalents. Hence, the inhibitory effect of BRICHOS against αSyn works under different environmental conditions, including near-physiological levels of salt (SI Fig. S1).

We further aimed to confirm our results by performing the same experiments at lower αSyn concentration, using 30 μM instead of 70 μM αSyn, and adding the same molar ratios of BRICHOS (SI Fig. S1). We indeed found that BRICHOS also delays αSyn aggregation at 30 μM (SI Fig. S1), suggesting a similar inhibitory mechanism, which is determined by the BRICHOS:αSyn ratio. Taken together, these results show that the inhibitory effect of BRICHOS is present at different αSyn concentrations and in the presence of near-physiological salt concentration.

### BRICHOS specifically inhibits secondary nucleation and fibril-end elongation

Amyloid formation can generally be described as nucleation-dependent polymerization mechanism (Cohen et al., 2012). The formation of a nucleus, a process known as primary nucleation (*k_n_*), triggers the growth of fibrils through fibril-end elongation (*k_+_*). Secondary processes can also occur such as monomer-dependent secondary nucleation (*k_2_*) or fragmentation (*k_−_*). αSyn aggregation kinetic traces can be fitted with these theoretical kinetics models, describing the contributions of each specific microscopic nucleation events (Buell et al., 2014; Gaspar et al., 2017; Knowles et al., 2009). Here, we applied a model consisting of three microscopic processes: primary nucleation, monomer-dependent secondary nucleation, and fibril-end elongation. In addition to monomer-dependent secondary nucleation, monomer-independent secondary nucleation reactions might be present, which are related to fibril fragmentation, and described by the nucleation rate constant *k_−_*. Typically, this process can be related to agitation during the fibril reaction (Cohen et al., 2013). Previous studies have shown that αSyn aggregation follows a secondary nucleation-dependent aggregation mechanism at physiological pH (Buell et al., 2014). By conducting a detailed global fit analysis of the aggregation kinetics using the webserver AmyloFit (Meisl et al., 2016), we could identify which nucleation rates are affected by BRICHOS (Fig. 1a).

First, we applied a model that only included monomer-dependent secondary nucleation reactions, in addition to primary nucleation and elongation. To decipher the contribution of the individual nucleation events, we set two nucleation rate constants as global fit parameters, meaning that they are constrained to the same value across all BRICHOS concentrations, while one nucleation rate constant was set as an individual fitting parameter that could vary with BRICHOS concentration (Fig. 1a). The global fits that best described the data were the ones where either *k_2_* or *k_+_* were free fitting parameters, suggesting that BRICHOS suppresses fibril secondary nucleation and/or elongation. The same results were obtained in the presence of 154 mM NaCl and at lower αSyn concentration (SI Fig. S1).

Second, we tested a different nucleation model where monomer-dependent secondary nucleation, *k_2_*, is neglected and replaced by fragmentation-related monomer-independent secondary nucleation, *k_−_*. Also, for this model we obtained overall good fits for either *k_−_* or *k_+_* as the free fitting parameters (SI Fig. S2), yet the mean residual errors, which show an increase by a relative factor of 5.5 for the fragmentation-dependent model, favor the monomer-dependent secondary nucleation model.

In order to deduce whether BRICHOS affects predominantly fibril elongation or secondary nucleation, or both, we performed highly seeded kinetics (Fig. 1c). At high seed concentration a large number of free fibril ends is available, creating a condition where the start of the aggregation kinetics is dominated by only fibril-end elongation (Cohen et al., 2012). Hence, we could extract the elongation rates by analyzing the slope of the first 2 h of the kinetic traces, which is proportional to the fibril-end elongation rate *k_+_* (Fig. 1c). We found that the initial slope exhibits a dependence on the BRICHOS concentration, indicating that BRICHOS prevents the elongation of αSyn fibrils (Fig. 1c,d). By comparing the dependencies of the combined fitting parameter *k_+_k_2_* from the global fit analysis (Fig. 1d) with the initial slope of the highly seeded aggregation kinetics, we found that the modulation of *k_+_* alone cannot explain the large effect observed on *k_+_k_2_*. This suggests that *k_2_* is predominantly reduced by the presence of BRICHOS. Hence, these results show that BRICHOS suppresses mainly secondary nucleation in addition to fibril-end elongation.

While fragmentation-related processes might be present as well, a model including both *k_−_* and *k_2_* exhibits several coupled fitting parameters, which increases the risk for over-fitting (Meisl et al., 2016). When we combine the slightly better fits for *k_2_*, with the fact that the inhibition of secondary nucleation on the fibrils surface can be mechanistically understood by bound BRICHOS on αSyn fibrils, we arrive at the simplest model for the mechanism-of-action for BRICHOS: it prevents mainly monomer-dependent secondary nucleation in addition to fibril-end elongation.

### BRICHOS exhibits an inhibitory effect on the aggregation of a C-terminally truncated αSyn variant and familial PD αSyn mutants

We then investigated the effect of BRICHOS on a more aggregation-prone C-terminally truncated αSyn variant as well as two prevalent familial αSyn mutants associated with PD (A30P and A53T). C-terminally truncated variants have been shown to exhibit an accelerated propensity for aggregation, where mainly secondary nucleation processes related to fragmentation are enhanced (Farzadfard et al., 2022). Here, we studied the effect of BRICHOS on a variant of αSyn C-terminally truncated at position 121, which is referred to as αSyn121. Our experiments revealed a modest but significant concentration-dependent inhibitory effect of BRICHOS on αSyn121 aggregation kinetics (Fig. 2a). Also in this case, a global fit analysis using a monomer-dependent secondary nucleation model identified secondary nucleation processes and/or fibril-end elongation to be suppressed by BRICHOS. Interestingly, we found that the reduction of the combined nucleation rate constant for secondary nucleation process, *k_+_k_2_*, is smaller than for WT αSyn (Fig. 2e). This can be rationalized by the differences in the nucleation mechanisms of αSyn121 and WT αSyn. A saturation of the elongation step and a substantially increased fragmentation rate was reported for αSyn121 (Farzadfard et al., 2022). BRICHOS fibril binding is unlikely to target fragmentation processes, resulting in an overall less efficient inhibitory effect of BRICHOS for αSyn121 aggregation. Additionally, different binding efficiency of BRICHOS to αSyn121 fibrils might contribute.

**Figure 2:**
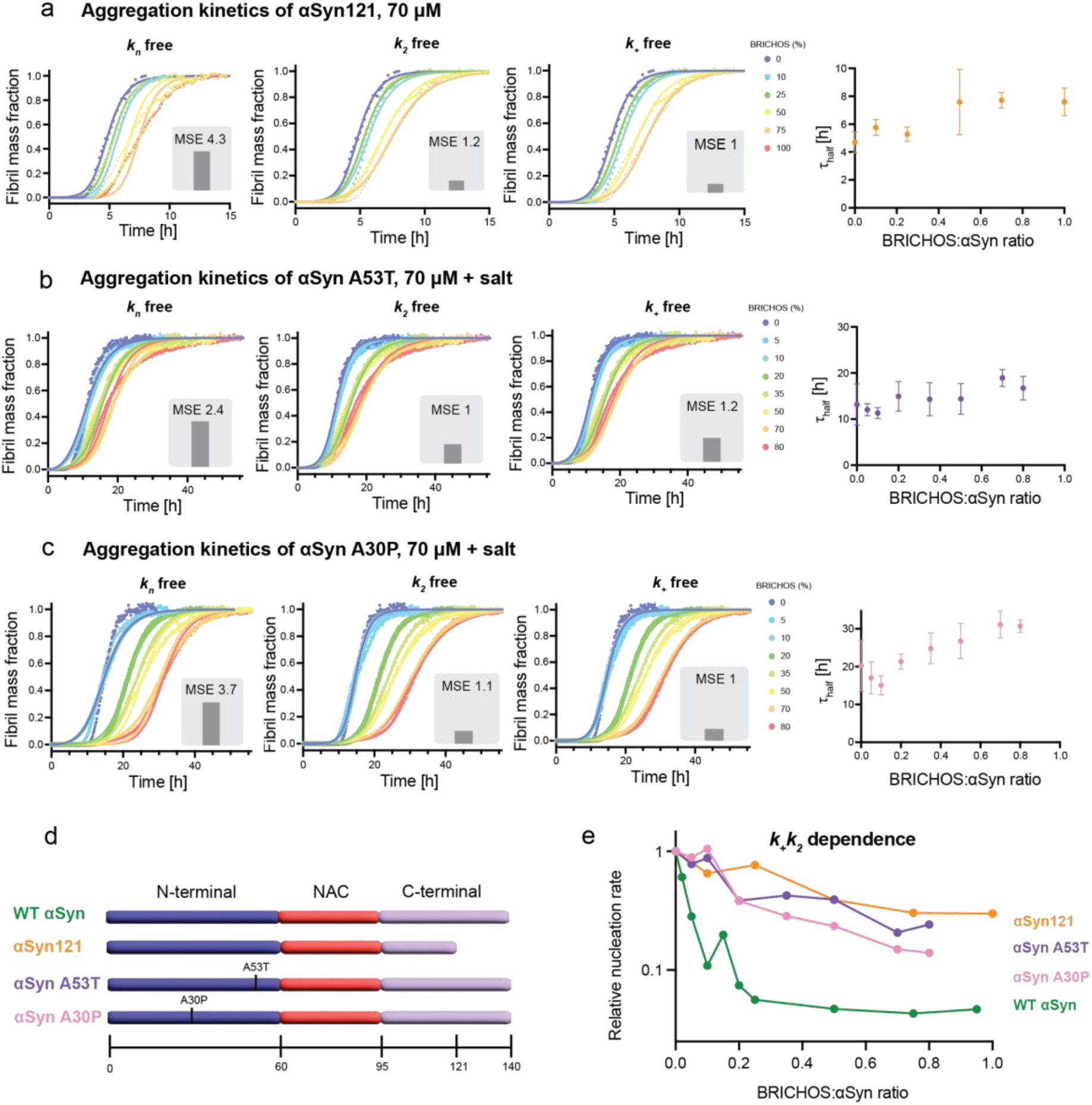
Inhibitory effect of BRICHOS on a C-terminally truncated αSyn variant and familial PD αSyn mutants. (a-c) ThT fluorescence aggregation kinetics of 70 µM (a) αSyn121, (b) A53T αSyn and (c) A30P αSyn in 20 mM NaP buffer, 0.2 mM EDTA at pH 7.4 (with 154 mM NaCl for A30P and A53T), using agitation and glass beads, with six replicates in the presence of 0 to 100 % BRICHOS (color gradient from blue to red) at 37 °C fitted to a secondary nucleation dominated kinetic model with Amylofit (Meisl et al., 2016). Individual fits (solid lines) to normalized and averaged aggregation traces (dots) of αSyn aggregation kinetics are shown in the presence of BRICHOS where either *k_n_*, *k_2_* or *k_+_* is the free fitting parameter. The normalized mean square error (MSE) value refers to the global fits. The aggregation half-time (τ_half_) as function of BRICHOS:αSyn ratio shows an inhibitory effect of BRICHOS on the aggregation of the αSyn variants. **(d)** Schematic view of WT αSyn, the C-terminally truncated variant αSyn121, and the familial PD mutants A53T and A30P, showing the N-terminal (purple), NAC (Non Amyloid Component) (red) and C-terminal parts (light purple) of the protein sequence. **(e)** Dependence of combined nucleation rate *k_+_k_2_* of the different αSyn variants on the BRICHOS:αSyn ratio, revealing a less pronounced inhibitory effect of BRICHOS on these αSyn variants compared to WT αSyn (without salt). The dependences of WT αSyn with and without salt are very similar and shown in SI Fig. S1.

Moreover, we tested the effect of BRICHOS on the familial αSyn mutants A30P and A53T found in familial PD. Under the same conditions, *i.e.* in presence of salt, the aggregation of A53T appears similar to WT αSyn with an aggregation half-time of 13.2 ± 4.5 h (Fig. 2b). In contrast, the aggregation half-time of A30P αSyn is longer with 20.2 ± 6.0 h (Fig. 2c). In the presence of increasing concentrations of BRICHOS, we found, similarly as for WT αSyn, a concentration-dependent effect where higher concentrations of BRICHOS increase the aggregation half-times (Fig. 2b,c). Also, for aggregation traces of the αSyn mutants, a nucleation model where only the secondary nucleation and/or fibril-end elongation rate are reduced, accurately fits the aggregation traces for both A30P and A53T kinetics (Fig 2b,c). Notably, the relative reduction in the combined nucleation constant *k_+_k_2_* is similar for both αSyn mutants, but considerably smaller than for WT αSyn (Fig. 2e). The A30P and A53T mutants were reported to exhibit different nucleation mechanisms compared to WT αSyn, with a larger contribution of primary nucleation (Ohgita et al., 2022), which may cause the reduced efficiency of BRICHOS in inhibiting aggregation.

### BRICHOS does not affect the morphology of αSyn fibrils

To investigate whether BRICHOS modulates the αSyn fibril morphology, we collected transmission electron microscopy images of αSyn fibrils at the end of the aggregation reaction, obtained without and with 100 % molar equivalent of BRICHOS (Fig. 3a,b). We measured 100 diameters of fibrils for both conditions and observed that the diameters of the BRICHOS co-incubated αSyn fibrils were 11.8 ± 2.4 nm compared to 12.3 ± 1.9 nm for αSyn fibrils alone (Fig. 3c). These values are in good agreement with the published *in vitro* structure of αSyn fibrils, whose diameter is around 10 nm corresponding to two αSyn molecules per layer (Guerrero-Ferreira et al., 2019; B. Li et al., 2018; Y. Li et al., 2018). In conclusion, BRICHOS does not have a major effect on the morphology of αSyn fibrils when present during aggregation.

**Figure 3:**
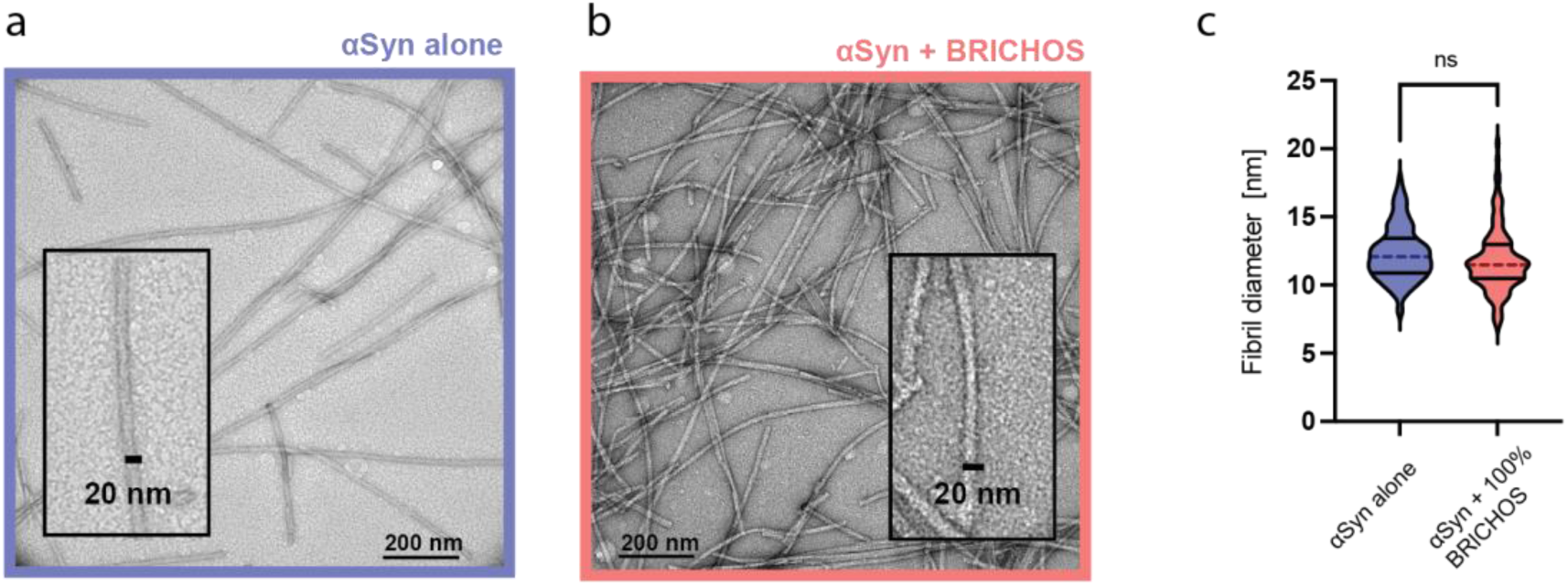
BRICHOS does not affect αSyn fibril morphology. **(a,b)** TEM images of αSyn fibrils formed (a) without or (b) with 100% BRICHOS molar equivalent. **(c)** Fibril diameter of 100 αSyn fibrils incubated without or with BRICHOS displayed as median (dashed lines) and quartiles (solid lines). An unpaired T-test revealed no significant difference in the fibril diameter between the two samples.

### BRICHOS binds to αSyn fibrils but not αSyn monomers

To gain further information on how BRICHOS is able to interfere with the αSyn fibrillation process, we employed various biophysical techniques, including Isothermal Titration Calorimetry (ITC), Surface Plasmon Resonance (SPR), Flow-induced Dispersion Analysis (FIDA) and solution Nuclear Magnetic Resonance (NMR), to assess the binding capacity of BRICHOS to αSyn monomers and fibrils. FIDA measures the diffusion of fluorescently labelled proteins in a microfluidic channel, which can be translated to a hydrodynamic radius *R*_h_. Binding to another (non-labeled) species leads to a change in *R*_h_.

First, the binding of BRICHOS to αSyn monomers was investigated. FIDA experiments revealed that the hydrodynamic radius of fluorescently labeled BRICHOS-Alexa^488^ is constant for the investigated αSyn monomer concentrations, indicating that BRICHOS does not bind to αSyn monomers (Fig. 4a). These results were confirmed by ITC, where titration of BRICHOS to αSyn did not result in any significant heat increase compared to the control experiment where BRICHOS was titrated on buffer alone (SI Fig. S3). Further, ^1^H-^15^N HSQC 2D NMR experiments were performed, where unlabeled BRICHOS was titrated onto ^15^N-labeled αSyn monomers. The observed signal loss was uniform along the protein sequence and corresponded to the dilution factor, confirming that BRICHOS does not bind to αSyn monomers (SI Fig. S3). Second, we studied the binding of BRICHOS to αSyn fibrils. In FIDA experiments, we detected high molecular weight species of BRICHOS-Alexa^488^ in the presence of fibrils, indicating binding to the fibrils (Fig. 4a); however, the shape of the binding curve and appearance of excessively large molecules prevented a precise determination of the binding constant. To solve this, BRICHOS-Alexa^488^ and the fibrils were mixed prior to the experiment allowing for estimations based on the shift in fluorescence intensity upon binding rather than the hydrodynamic radius, which pointed towards a K_D_ value of 2.3 ± 0.4 μM in terms of αSyn monomer equivalents (SI Fig. S3). These findings were supported by SPR experiments using unlabeled proteins, where αSyn fibrils were immobilized on the SPR chip. The SPR binding curves exhibited BRICHOS binding to αSyn fibrils. SPR traces could be fitted with a two-phase binding model (Fig. 4b). This analysis revealed apparent dissociation constants for one weak and one strong binding of 350 ± 60 µM and 22 ± 2.0 nM, respectively. The rough estimate of the K_D_ from the FIDA experiments hence represents a mix of these two binding processes observed by SPR.

**Figure 4:**
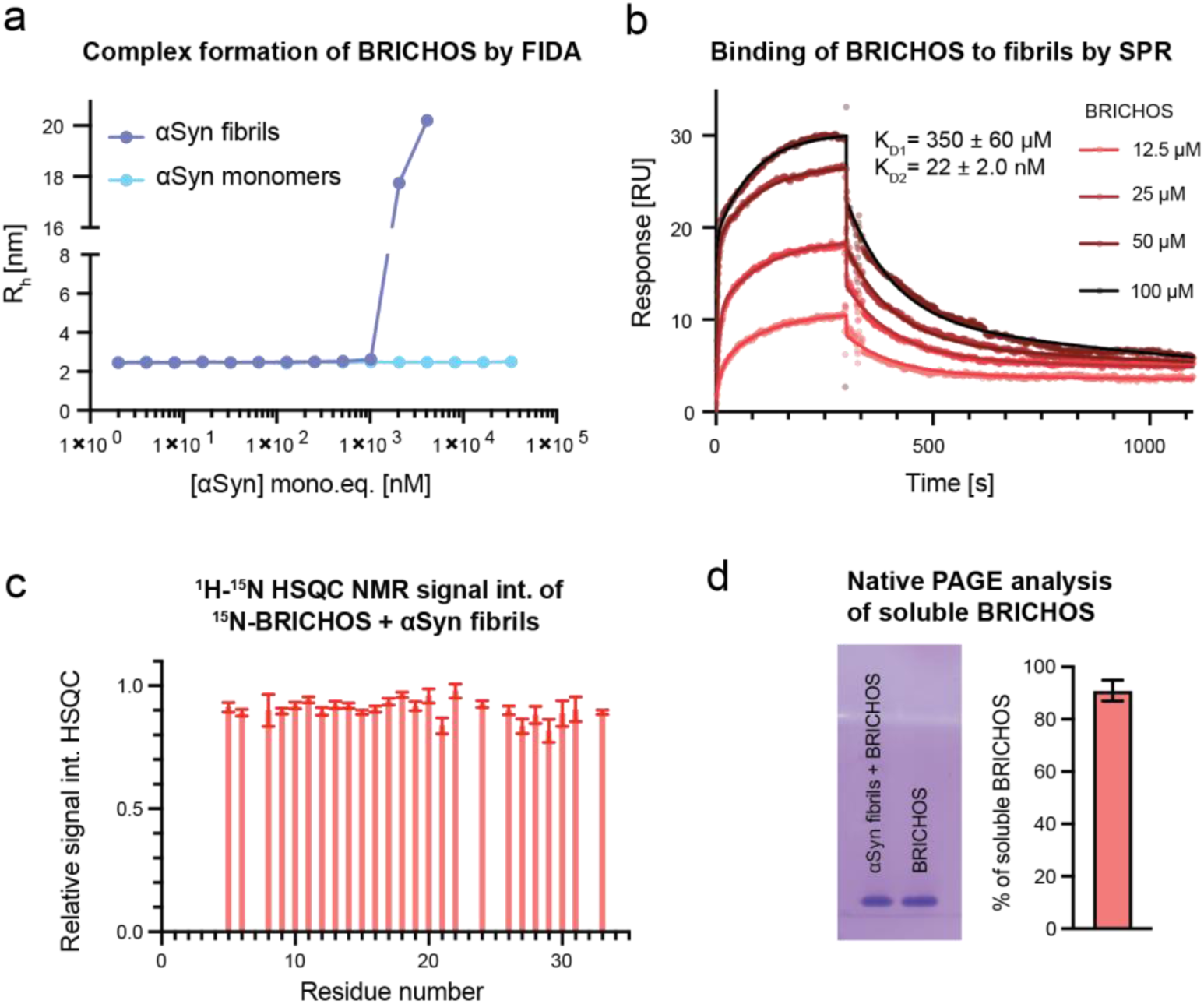
BRICHOS binds to αSyn fibrils but not to αSyn monomers. (a) FIDA experiments measuring the hydrodynamic radius (R_h_) of 50 nM fluorescently labeled BRICHOS-Alexa^488^ in the presence of WT αSyn monomers or WT αSyn fibrils (in monomer concentration equivalents), showing no complex formation of BRICHOS with αSyn monomers but binding to αSyn fibrils. **(b)** SPR measurements of BRICHOS binding to αSyn fibrils in 20 mM sodium phosphate, 0.2 mM EDTA, pH 7.4, revealing a two-phase profile with a weak and strong dissociation constant of 350 ± 60 µM and 22 ± 2.0 nM, respectively. **(c)** Relative ^1^H-^15^N HSQC NMR signal intensities of ^15^N-labeled BRICHOS in the presence of equimolar ratio of sonicated αSyn fibrils (referred to the monomer equivalent concentration), showing a relative average signal of 90.5 ± 3.8 % compared to without the presence of sonicated αSyn fibrils. **(d)** Native PAGE analysis of soluble BRICHOS in the presence of αSyn fibrils, exhibiting 90.6 ± 4.2 % of soluble, unbound BRICHOS. The uncropped gel is shown in SI Fig. S3.

To determine the amount of BRICHOS bound to αSyn fibrils, we performed solution NMR experiments recording ^1^H-^15^N HSQC spectra, where sonicated unlabeled αSyn fibrils were added to ^15^N-labeled BRICHOS. Due to NMR-unfavorable exchange dynamics of BRICHOS (Leppert et al., 2023), only parts of the expected resonances are visible in the NMR spectra. Bound BRICHOS is expected to be invisible in the ^1^H-^15^N HSQC spectrum due to the large size of αSyn fibrils. We found that addition of sonicated αSyn fibrils at equimolar concentration resulted in a uniform attenuation of NMR signal to 90.5 ± 3.8 %, indicating that only a small proportion of BRICHOS is bound (Fig. 4c and SI Fig. S3). Further, the soluble fraction of BRICHOS in co-incubated BRICHOS-αSyn fibril samples was estimated by native PAGE gel analysis. Comparing BRICHOS alone to the co-incubated samples showed a reduction of the band intensity to 90.6 ± 4.2% (Fig. 4d and SI Fig. S3). Hence, both NMR and Native PAGE experiments support each other, suggesting that ∼ 10 % of BRICHOS is bound to αSyn fibrils under the conditions tested.

### BRICHOS prevents αSyn oligomer generation by binding to αSyn fibrils and oligomers

The preference of BRICHOS binding towards structured αSyn fibrils compared to unstructured αSyn monomers raised the question whether BRICHOS can also target partly structured αSyn complexes, such as structures formed before the fibrillar state is reached. Partly structured αSyn complexes here refer to oligomeric or low molecular weight fibrillar species. We started first by investigating the ability of BRICHOS to modulate the rate of oligomer generation. From the detailed analysis of aggregation kinetics, the rate of generation of new nucleation units can be calculated from the contributions of the individual microscopic nucleation rate constants (Cohen et al., 2015). Since these nucleation units subsequently convert into oligomeric or low molecular weight fibrils, the nucleation rate gives an estimate about the rate of formation of new intermediate oligomeric/fibrillar species. Applying this model, the nucleation rates can be determined for different BRICHOS concentrations (Fig. 5a). We found that the maximum of the reaction is shifted to longer time points and that the area under the curve, which refers to the number of new nucleation units formed, is substantially decreased with increasing BRICHOS concentration. Hence, based on the estimations from the kinetic analysis, BRICHOS efficiently reduces the generation of new oligomeric or low molecular weight species (Fig. 5b).

**Figure 5:**
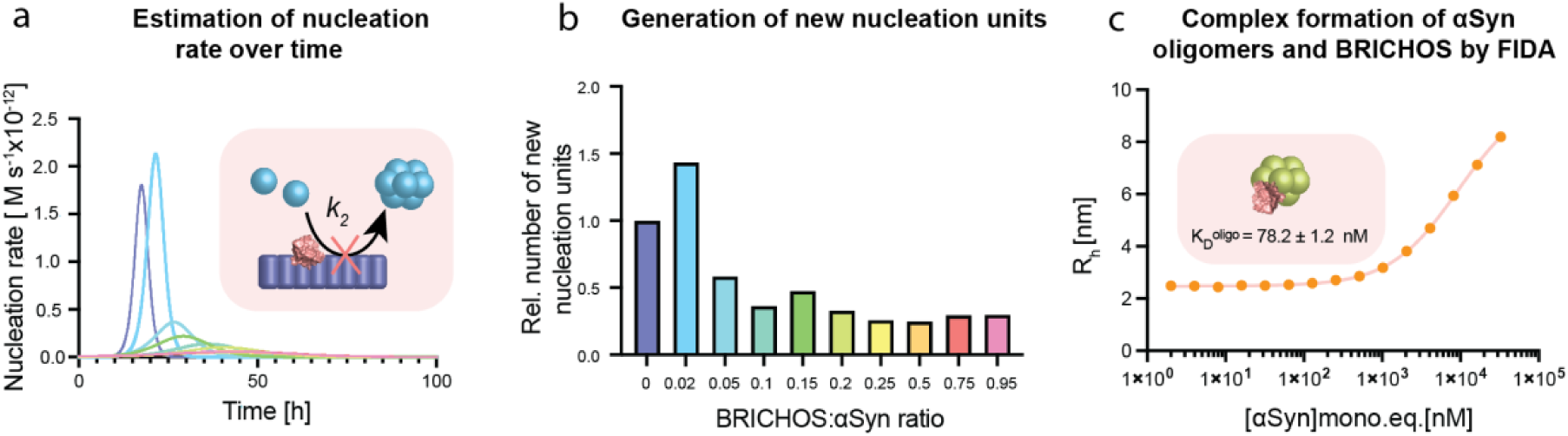
BRICHOS binds to αSyn oligomers and reduces their generation by inhibiting secondary nucleation. (a) Rate of formation of new nucleation units from global fit analysis of WT αSyn aggregation in the presence of different BRICHOS:αSyn ratios (blue to red color gradient), which is mainly determined by the reduction of secondary nucleation processes by BRICHOS. **(b)** Estimation of the number of new nucleation units at different BRICHOS:αSyn ratios, showing a substantial decrease in the presence of BRICHOS. **(c)** FIDA experiments of hydrodynamic radius (R_h_) of 50 nM BRICHOS-Alexa^488^ in presence of αSyn oligomers (in monomer equivalents), revealing a binding constant of 78.2 ± 1.2 nM.

Of note, various oligomeric forms can emerge during the aggregation process and their different characteristics and diverse structural arrangements greatly depend on the experimental conditions used. We therefore investigated whether BRICHOS, besides binding to αSyn fibrils and modulating the rate of oligomer formation, also directly binds to oligomeric αSyn species. We prepared αSyn oligomers by incubating αSyn monomers at 37 °C during 3 h with shaking at 900 rpm, followed by the isolation of the oligomeric species through size exclusion chromatography after the removal of larger aggregates by centrifugation (Paslawski et al., 2016). These αSyn oligomers of ∼ 30 molecules of αSyn are highly stable, kinetically trapped, show β-structure content, and are able to permeabilize lipid vesicles *in vitro* (Lorenzen et al., 2014). By performing FIDA experiments, we observed that fluorescently labeled BRICHOS-Alexa^488^ can bind to these αSyn oligomers with an apparent dissociation constant of K_D_ ^oligo^ = 78.2 ± 1.2 nM (Fig. 5c).

In conclusion, these findings demonstrate that, besides binding to the fibril surface and interfering with secondary nucleation, BRICHOS directly targets β-sheet containing αSyn oligomers.

### BRICHOS suppresses αSyn-associated neurotoxicity to hippocampal electrophysiology

Finally, we assessed how the impact of BRICHOS on αSyn aggregation translated into modulation of αSyn-related neurotoxicity. Electrophysiological assays were conducted on hippocampal brain slices from WT mice. *γ*-Oscillations in the brain are closely intertwined with cognition, learning and memory, and they have been reported to be impacted in dementia patients suffering from cognitive decline (Arroyo-García et al., 2021; Schnitzler & Gross, 2005). This makes *γ*-oscillations convenient indicators of neuronal activity and the neurotoxicity induced by external compounds (Kurudenkandy et al., 2014). Here, we used short αSyn fibrils, obtained after sonication, to induce neurotoxicity. After exposure to control buffer (PBS with 0.1 % sodium azide), sonicated αSyn fibrils alone or in combination with equimolar concentrations of BRICHOS were added to the hippocampal slices and incubated for 30 minutes. Subsequently, *ex vivo γ*-oscillations were elicited with kainic acid (KA; 100 nM) and recorded (Fig. 6). 500 nM sonicated αSyn fibrils (calculated from initial monomer concentration) did not result in any significant reduction in *γ*-oscillation power, while a concentration of 1 μM sonicated αSyn fibrils led to a significant (p < 0.001) decrease in *γ*-oscillation power, indicating neurotoxic effects. Importantly, co-incubation of 1 μM sonicated αSyn fibrils with BRICHOS at equimolar concentrations resulted in no impact on the power of *γ*-oscillations, suggesting that the presence of BRICHOS effectively blocked the neurotoxic effects of the sonicated αSyn fibrils.

**Figure 6:**
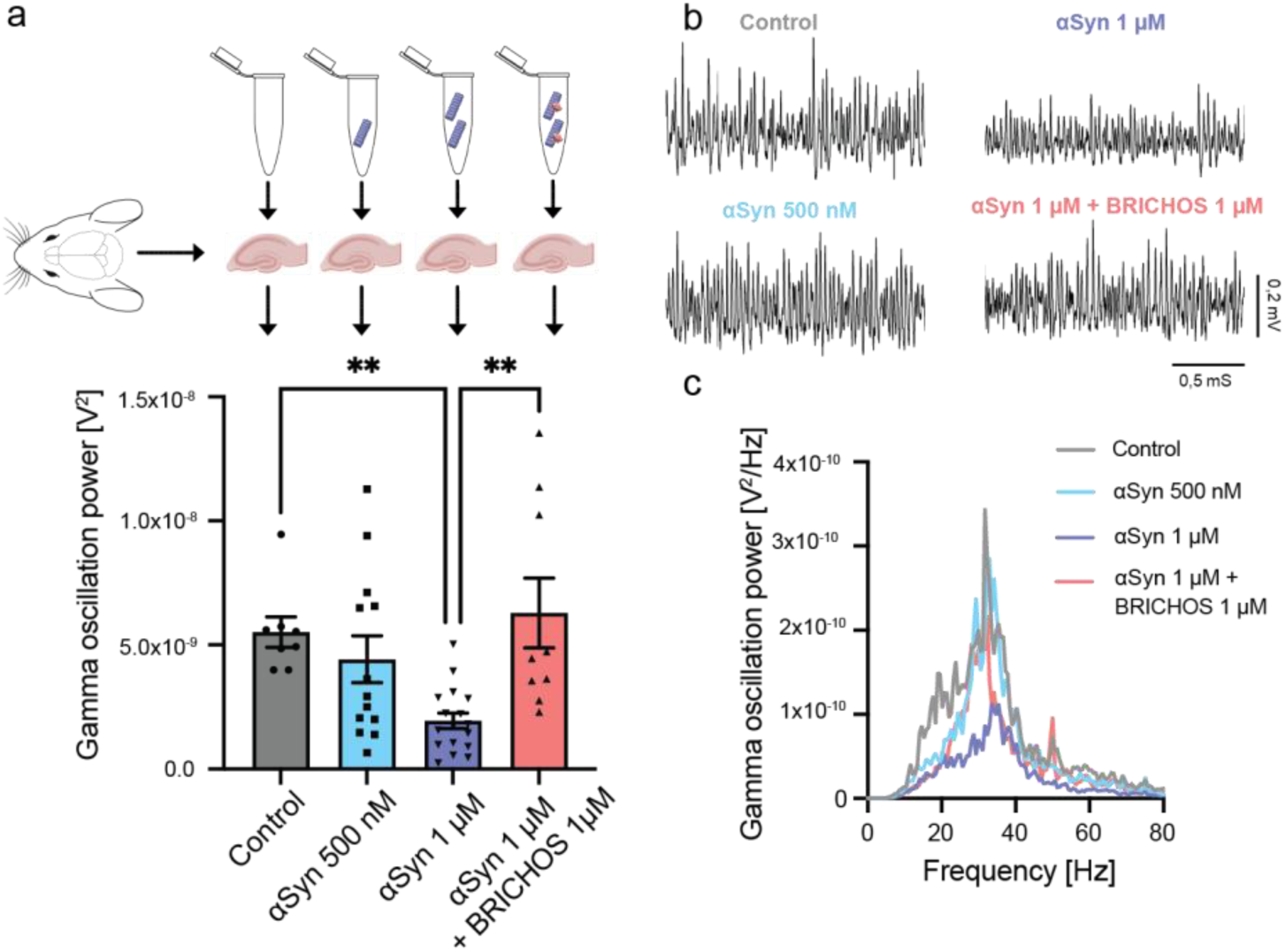
BRICHOS prevents neurotoxic effects of sonicated αSyn fibrils in hippocampal electrophysiology. (a) Schematic experimental set-up where the toxicity of sonicated WT αSyn fibrils with and without BRICHOS is measured as the impact on γ-oscillation power of hippocampal mouse brain slices. **(b)** Examples of *γ*-oscillation power spectra. **(c)** Example traces of *γ*-oscillations under control condition (gray), after incubation with 500 nM (light blue) or 1 μM (blue) sonicated αSyn fibrils, or 1 μM of sonicated αSyn fibrils with equimolar concentration of BRICHOS (red). One-way ANOVA revealed that sonicated αSyn fibrils at 1 μM exhibited significant (p < 0.001) toxic effects, which are efficiently prevented by the presence of BRICHOS.

Sonicated fibrils can release monomeric and oligomeric αSyn, which were shown to display toxic effects in neuronal cells (Cascella et al., 2021). The observed inhibitory effect of BRICHOS observed here could hence be explained by the fact that BRICHOS can prevent secondary nucleation-mediated generation of new nucleation units and/or directly target oligomeric αSyn species, which were either released from the fibrils and/or originated from endogenous αSyn.

## DISCUSSION

In this study, we show that a molecular chaperone domain can be utilized as an efficient inhibitor of secondary nucleation processes by binding to the αSyn fibrils surface and to αSyn oligomers, preventing the generation of oligomeric and low molecular weight fibrillar species as well as αSyn-associated toxicity (Fig. 7). Interestingly, a similar inhibitory mechanism was reported for BRICHOS on the aggregation and toxicity of different Aβ variants, including Aβ42, Aβ40 and artic E22G Aβ (Chen et al., 2017; Chen et al., 2020; Poska et al., 2016; Zhong et al., 2022). Also, in the case of Aβ, the binding of the BRICHOS domain (here a monomer-stabilizing mutant of the BRICHOS domain from lung surfactant protein C precursor, referred to as proSP-C BRICHOS), to small, ThT-active and secondary nucleation competent fibrillar aggregates, consisting of ≤ 8 monomers, was shown (Leppert et al., 2021). These aggregates are secondary nucleation competent (Leppert et al., 2021), indicating that BRICHOS has the capacity to recognize secondary nucleation sites already at early aggregation state intermediates, and our results indicate that similar mechanisms appear to be present for αSyn. Another study determined the efficiency of proSP-C BRICHOS in reducing the Aβ oligomer formation rate, where the oligomers were found to be mainly on-pathway (Linse et al., 2020; Michaels et al., 2020). Furthermore, recent treatment studies of AD mouse models using BRICHOS exhibited a positive effect on neuroinflammation and cognitive behavior (Manchanda et al., 2022), which could be correlated to the inhibition of oligomer generation *in vitro* (Abelein & Johansson, 2023). The findings from these investigations, coupled with the outcomes of the present study, collectively demonstrate the efficacy of BRICHOS in targeting secondary nucleation and preventing the formation of oligomeric species, which seemingly protects against toxicity.

**Figure 7:**
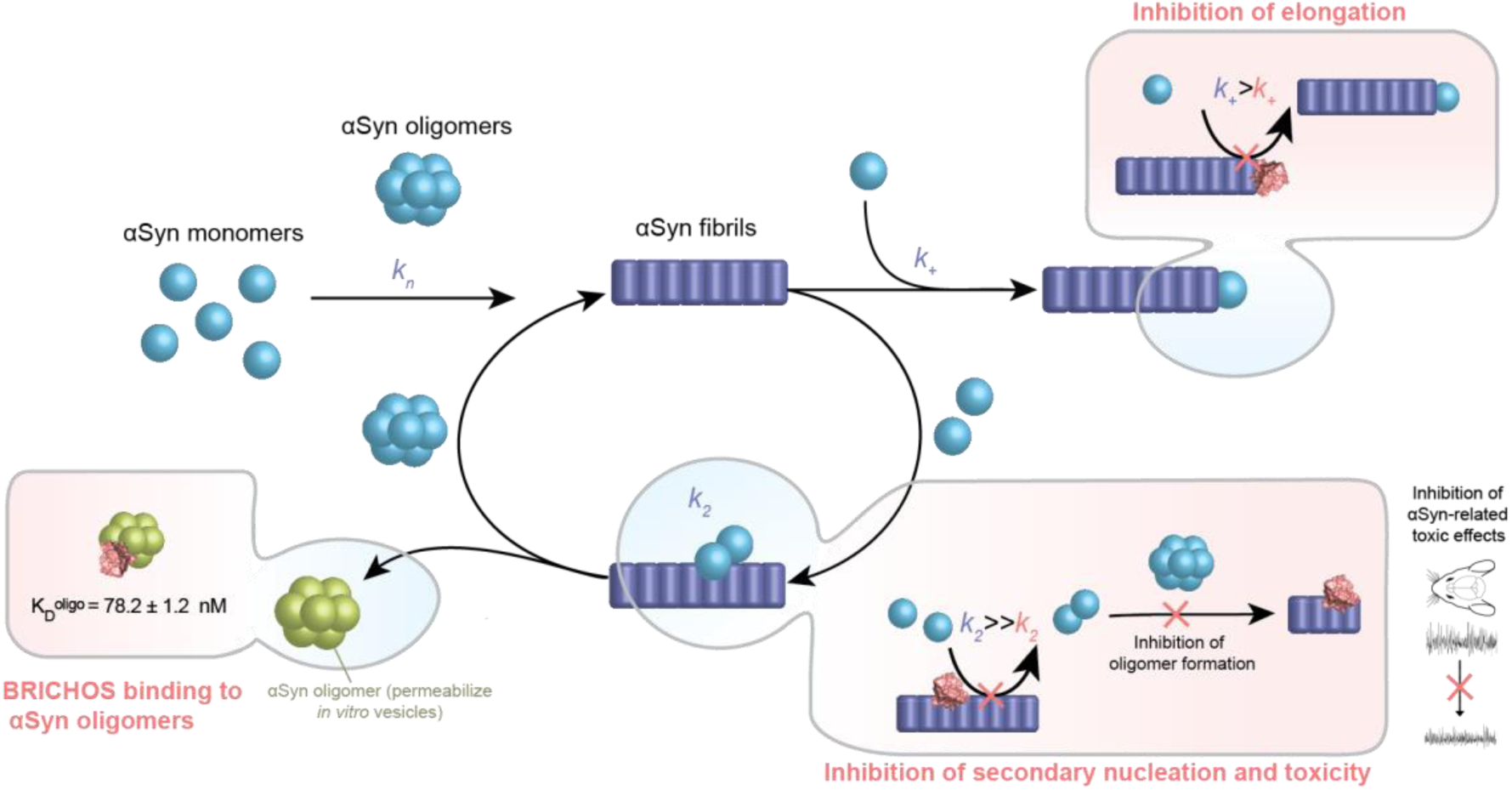
Model for mechanism-of-action of inhibition of αSyn aggregation by the molecular chaperone domain BRICHOS. BRICHOS (depicted in light red as 3D model from I-Tasser) has the capacity to inhibit fibril-end elongation (*k_+_*) and predominantly secondary nucleation (*k_2_*) by binding to the αSyn fibril surface, which reduces the generation of new nucleation units. These nucleation units can convert into oligomeric or low molecular weight fibrillar species, generating an autocatalytic process, which is blocked by BRICHOS. Furthermore, besides targeting fibril surfaces, BRICHOS binds specific oligomeric αSyn species (green), which were reported to permeabilize vesicles *in vitro* (Lorenzen et al., 2014). Hence, BRICHOS suppresses secondary nucleation processes and secondary nucleation products, thereby exerting neuroprotective effects against the toxicity associated with αSyn.

The apparent generic ability of BRICHOS to bind to the fibril surfaces of various amyloid fibrils (Biverstål et al., 2020) points toward a common mechanism of BRICHOS recognition of amyloid fibrils, as demonstrated for Aβ and αSyn. One hypothetical mechanism is that BRICHOS targets specific sites on the fibril surface, which would otherwise act as catalyzers for secondary nucleation. Importantly, also small oligomeric species, which cannot convert into fibrils can be bound by BRICHOS, further diminishing the spread of these potentially neurotoxic species. Such off-pathway oligomers have also been produced without shaking and characterized to induce cellular toxicity by disrupting cell membrane in neuroblastoma cells and rat primary cortical neurons (Fusco et al., 2017) and permeabilize cell membrane model *in vitro* (Chen et al., 2015). Dual targeting of catalytic sites for secondary nucleation as well as secondary nucleation products by BRICHOS may explain the drastically remediated toxic effects associated with Aβ or αSyn aggregation and aggregation products. Future investigations evaluating the toxic effects of different αSyn species, such as αSyn oligomers and different fibrillar αSyn species, and the impact of BRICHOS will help to further elucidate how oligomer binding of BRICHOS translates into modulated neurotoxic effects.

Taken together, data in this report show that developing therapeutics against catalytic sites on fibril surface that mediate secondary nucleation as well as against secondary nucleation products, such as oligomeric or low molecular weight fibrillar species, may efficiently prevent amyloid-associated toxic effects. These mechanistic insights presented here for a molecular chaperone may be valuable for the design of other specific therapeutics, also including small molecule drugs and antibodies, with potential against α-synucleopathies. Our results also encourage the evaluation of molecular chaperone-based therapeutic approaches against αSyn-related neurodegenerative disorders *in vivo*.

## MATERIALS AND METHODS

### Preparation of Bri2 BRICHOS monomers

Recombinant R221E Bri2 BRICHOS monomers were produced in-house using adapted protocols as published previously (Chen et al., 2020). In detail, SHuffle T7 (K12 strain) *E. coli* cells were transformed with a plasmid vector carrying the human Bri2 BRICHOS R221E mutant sequence (113–231 of full-length human Bri2) fused to Gly-His6x-NT* at the N-terminus separated by a thrombin cleavage site. The cells were incubated at 30 °C at 120 rpm in LB medium containing 15 μg/ml kanamycin until reaching an OD of 0.8. The cells were then induced with 0.5 mM IPTG and grown overnight (O/N) at 20 °C. The cells were harvested at 5,000 x g centrifugation at 4 °C and pellets were resuspended into 20 mM Tris HCl, pH 8.0 and put at −20°C. Two 1L pellets were thawed and one tablet of protease inhibitor (cOmplete^™^, Roche) was added. Each pellet was sonicated for 5 min on ice (2 s ON, 2 s OFF, 65 % of max power) (Vibra-Cell™ model CV334, SONICS) and the resulting lysate centrifuged at 24,000 x g, 4 ℃ for 30 min. The collected supernatant was then loaded on a pre-equilibrated Ni-NTA column (GE healthcare Bio-Science AB, Sweden). The column was subsequently washed with 30 column volumes (CV) using 20 mM Tris HCl, 15 mM imidazole, pH 8.0 followed by elution using 20 mM Tris HCl, 300 mM imidazole, pH 8.0. The fractions containing BRICHOS were pooled and dialyzed overnight against 20 mM sodium phosphate, 0.2 mM EDTA, pH 7.4 at 4℃. To cleave off the N-terminal NT* tag, His-tagged thrombin was added during the dialysis (1:1000 enzyme to substrate, w/w). The sample was then loaded on Ni-NTA gravity columns to separate His-NT* and His-thrombin from the protein of interest. The flowthrough was concentrated using 10 kDa cut-off concentrator (Vivaspin 20, 10 kDa MWCO Cytiva), heat treated in a water bath for 10 min at 60 °C and cooled down before loaded on a Hiload Superdex 75 PG Size Exclusion column (Cytiva, USA) to collect the peak of interest corresponding to the monomeric BRICHOS. The concentration of the protein sample was measured using the optical density at 280 nm using the extinction coefficient of 9065.

### Preparation of αSyn monomers and oligomers

Recombinant αSyn121 was purified according to published protocol until the lyophilization step (Farzadfard et al., 2022; Lorenzen et al., 2014; Paslawski et al., 2016) and further purified using size exclusion chromatography as mentioned below. Recombinant WT, A53T and A30P αSyn monomers were obtained as described in the literature (Kumar et al., 2020) applying slightly adopted purification steps. In this study, BL21(DE3) *E.coli* were transformed with the bacterial expression plasmid pET21a-αSynuclein (Addgene) containing the human αSyn coding sequence. For both A53T and A30P αSyn familial PD mutants, site-directed mutagenesis was used using the vector pET21a-αSynuclein with the kit QuikChange XL Site-Directed Mutagenesis (Agilent technologies, USA) and the inserted mutations were checked by sequencing. To produce WT, A53T and A30P αSyn, the cells were grown at 37 °C in LB medium containing 100 μg/ml Ampicillin until reaching OD=0.8. The culture was inoculated with 0.5 mM IPTG and incubated at 20 °C O/N. The cells were centrifuged at 5,000 x g for 20 min at 4 °C and the bacterial pellets were resuspended by gentle vortexing in 20 mM Tris-HCl, pH 8.0 and kept at −20 °C. A cell pellet from 1 l was thawed and lysed using a sonicator (Vibra-Cell™ model CV334, SONICS) at 65 % maximum power for 5 min on ice with 2 s ON, 2 s OFF pulse setting. The sample was then boiled at 100 °C for either 5 or 10 min. This was followed by centrifugation at 24,000 x g for 20 minutes at 4 °C to remove degraded proteins. The supernatant, with the heat-stable αSyn protein, was loaded onto a HiTrap QFF anion exchange chromatography column. The protein was eluted with a gradient of 20 mM Tris HCl, 1 M NaCl, pH 8 during 10 CV. The eluted peak was then loaded on a reverse phase chromatography column using 99.9% H_2_O, 0.1% TFA as the running buffer. Elution was done using 70% ACN, 0.1% TFA buffer gradient during 20 CV. The fractions corresponding to the peak of interest were pooled and the concentration was measured using the optical density at 280 nm using the extinction coefficient of 5960. The samples were lyophilized O/N using a vacuum (*LabConco CentriVap Concentrator*). The lyophilized samples that were not used directly were stored at −20°C. For further purification, around 8 mg of lyophilized protein aliquots (also applied for αSyn121) were dissolved in 550 μl 7 M Gdn-HCl, pH 8.0 and loaded onto the Superose 6 Increase column with 20 mM sodium phosphate, 0.2 mM EDTA, pH 7.4 as running buffer to obtain homogenous and pure αSyn monomeric sample further stored at −20 °C until use.

Oligomeric αSyn were prepared as previously described (Farzadfard et al., 2022; Lorenzen et al., 2014; Paslawski et al., 2016).

### Thioflavin T (ThT) fluorescence aggregation assays

For ThT kinetics experiments, each sample contained either 30 or 70 μM of WT, αSyn121, A53T and A30P αSyn monomers in the presence of 20 μM ThT and different concentration of BRICHOS in 20 mM sodium phosphate, 0.2 mM EDTA, pH 7.4 buffer. For salted conditions, the same buffer with 154 mM NaCl was used. For A30P and A53T only, 0.02% of NaN_3_ in the buffer was used. Two glass beads of 1 mm diameter were used in each well. The fluorescence was recorded using a plate reader (FLUOStar Omega from BMG Labtech, Offenberg, Germany) and a 448-10 nm excitation filter and a 482-10 nm emission filter while the plate was orbitally shaken between cycles at 300 rpm for 9 min on and 1 min off. Each cycle was set to 10 min long and the temperature was set to 37 °C. For the preparation of fibril seeds, αSyn fibrils were obtained from previous runs of ThT kinetics using 70 μM αSyn monomers alone. These fibrils were then sonicated for 3 minutes, 2 s ON, 2 s OFF, 20% Amplitude using a probe sonicator (Vibra-Cell™ model CV18, SONICS) and 5 % of seeds were used.

### Analysis of αSyn aggregation kinetics

Aggregation traces were normalized, averaged over six replicates, and truncated at the plateau of the aggregation reaction. The aggregation kinetics were fitted by applying the secondary nucleation dominated, seeded nucleation model in the online platform AmyloFit (Meisl et al., 2016), which includes the rate constants for primary and secondary nucleation as well as fibril-end elongation. To evaluate the contribution of a single nucleation rate, two nucleation rate constants were set as global fit parameters, meaning that they are constrained to the same value across all BRICHOS concentrations, while one nucleation constant (*k_n,_ k_2,_* or *k_+_*) was set as an individual fitting parameter. MSE values were collected and normalized for each set of fitting. The *k_+_k_2_* dependencies on BRICHOS concentration were obtained using the unseeded secondary nucleation dominated model by constraining the combined rate constant *k_n_k_+_* as global fit constant. Highly seeded aggregation kinetics were analyzed using GraphPad Prism 10, where elongation rates were extracted by fitting a straight line to the first 2 h of the aggregation curves.

### Transmission electron microscopy

αSyn monomers (70 μM) without or with BRICHOS (molar ratio 1:1) were incubated similarly as for the ThT kinetic experiment but without ThT. After reaching around 100 h of experiment, the fibrils were spun down at 16,200 x g at 4 °C for 30 min and washed two times with fresh buffer. A 5 µl diluted solution of αSyn fibrils (around 10 µM) was applied on 200 mesh formvar coated nickel grid and excess solution was removed using blotting paper after 10 min of incubation. It was then washed twice with 10 µl MQ water and subsequently stained with 1% uranyl formate for 5 min. Excess stain was removed with blotting paper and the sample was subsequently air-dried. Transmission electron microscopy (FEI Tecnai 12 Spirit BioTWIN, operated at 100 kV) was performed for analysis of fibril morphology using 2 k × 2 k Veleta CCD camera (Olympus Soft Imaging Solutions, GmbH, Münster, Germany). 15 to 20 images were recorded for each sample randomly and fibril diameters were measured using the Image J software using 100 fibrils in each condition coming from different images. For long fibrils, 2 to 3 measurements were made along the fibril axis. An unpaired t-test was performed to compare the two samples.

### Native PAGE

BRICHOS was added to an equimolar concentration to αSyn fibrils sample (8 μM), previously prepared by incubating monomers (around 200 µM) in an Eppendorf tube at 37°C on a shaking incubator at 1000 rpm for 4 days. Samples were prepared under non-denaturing conditions in triplicates and subsequently run using Native gel at 7.5 %. BRICHOS was visualized using Coomassie blue staining.

### Surface Plasmon Resonance (SPR)

The binding analysis was performed using Biacore 3000 instrument at 25 °C. Different Chips were used to perform these experiments using αSyn fibrils, αSyn fibrils produced in the presence of salt and αSyn monomers. The system was primed with 20 mM sodium phosphate, 0.2mM EDTA, pH 7.4. The different concentration of BRICHOS (0, 1.56, 3.125, 6.25, 12.5, 25, 50 and 100 μM) were injected for 5 minutes at a flow rate of 20 μl/min over the channels 2 and 4. Channels 1 and 3 were left empty and served as a reference to control for possible non-specific binding of αSyn. Each sample was run in duplicates. The analysis consisted of first adjusting the sensorgrams to zero and removing buffer spikes. The signal of the three lowest BRICHOS concentrations used was small and therefore not included in the analysis. The curves were then fitted with a two-phase association and dissociation model using GraphPad Prism with the association rate determined by *R*_*Asso*_(*t*) and the dissociation rate by *R*_*Diss*_(*t*).

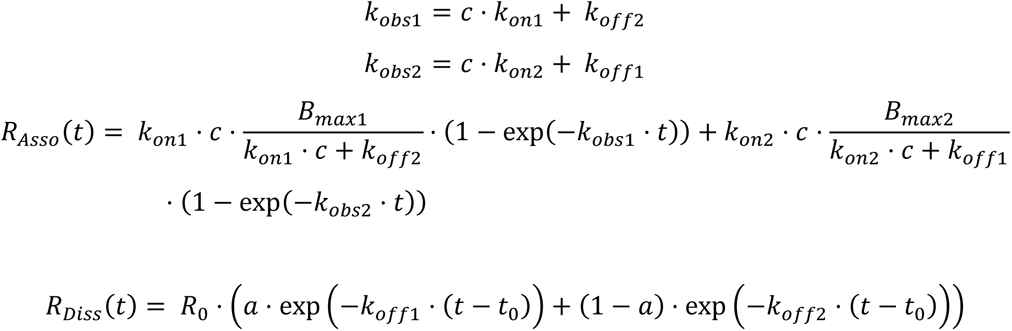

where *k*_*on*_ and *k_off_* refer to the on- and off-rate constants, *k*_*obs*1_ and *k*_*obs*2_ are the observed association rate constants, *c* is the BRICHOS concentration, *B*_*max*1_ & *B*_*max*2_ and *R*_0_ are the amplitudes for association and dissociation, respectively, and *a* is a value between 0 and 1, describing the contribution of the two dissociation phases. The fast and slow apparent K_D_ values were calculated as a ratio of the corresponding dissociation rate constants and association rate constants. The given errors correspond to the fitting error.

### NMR experiments

2D NMR ^1^H-^15^N HSQC experiments were performed using a 700 MHz Bruker Avance NMR spectrometer equipped with a triple resonance cryogenic probe. Titration measurements were conducted with ^15^N-labeled αSyn or ^15^N-labeled BRICHOS in 20 mM NaP, 0.2 mM EDTA with 10% D_2_O at pH 7.4 with additions of either BRICHOS or sonicated αSyn fibrils, respectively, at 281 K or 298 K, respectively. The peak assignment for αSyn was transferred from BMRB entry 18857 at pH 7.34. All NMR data were processed using Bruker Topspin 4.3 and NMRPipe (Delaglio et al., 1995), and analyzed with NMRFAM Sparky (Lee et al., 2009).

### Isothermal Titration Calorimetry (ITC)

216 μM BRICHOS monomers was injected 18 times with the injection syringe of 2 μl in the sample cell containing 20 μM αSyn monomers or buffer (20 mM NaP, 0.2 mM EDTA, pH 7.4) as the control using an iTC200 αSyn Microcal, GE Healthcare. The experiment was performed at room temperature with a stirring speed of 1,000 rpm.

### Flow-Induced Dispersion Analysis (FIDA)

BRICHOS was labeled with Alexa-488 by mixing 43.6 µM BRICHOS dissolved in 100 mM bicarbonate buffer (pH 8.3) and Alexa-488 NHS Ester (Thermo Scientific) dissolved in DMSO in a 1:2 molar ratio followed by 1 h incubation at RT. The labeled BRICHOS was separated from unreacted label by a PD-10 column (GE Healthcare) equilibrated with PBS. Flow-induced dispersion analysis was performed on a FIDA One instrument (FIDA Biosystems) using fluorescence detection with excitation wavelength at 480 nm. Standard 75 μm capillary (FIDA Biosystems) was rinsed by 1 M NaOH and equilibrated with PBS at 25 °C. The experiments were performed by priming the capillary by running buffer at a pressure of 3500 mbar. Subsequently, the analyte (αSyn) at various concentrations and indicator (BRICHOS-Alexa^488^) at 50 nM were sequentially injected into the capillary using the standard FIDA injection cycle (Pedersen et al., 2019). The resulting Taylograms were analyzed using FIDA software V2.3 (FIDA Biosystems), which provides hydrodynamic radius (R_h_) for each binding reaction. The resulting isothermal binding curves were fit to the following equation:

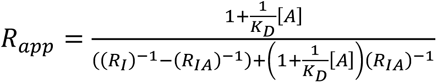

where K_D_ is the dissociation constant, [A] is the analyte concentration, and R_I_ and R_IA_ are the R_h_ of the indicator and the complex, respectively. Analysis of binding to the fibrillated αSyn species was based on shift in the BRICHOS-Alexa488 fluorescence signal upon titration, where indicator and analyte were allowed to mix prior to the injection. Here, the bound fraction (f_bound_) was calculated by normalizing the RFU signal to the highest and lowest values measured. The resulting dataset was fit the following isotherm equation:

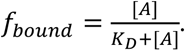

### Electrophysiological studies

For electrophysiological experiments, all chemical compounds used in extracellular solutions were obtained from Sigma-Aldrich Sweden AB (Stockholm, Sweden). Kainic acid (KA) was obtained from Tocris Bioscience (Bristol, UK). αSyn fibrils were prepared by incubating αSyn monomers at 37 °C at 900 rpm in PBS, 0.1% sodium azide, pH 7.4 for 7 days. They were then sonicated during 3 min, 2 s ON, 2 s OFF, 20% Amplitude using a probe sonicator (Vibra-Cell™ model CV18, SONICS).

We used WT mice (n=12) at 4-6 postnatal weeks (ethical permit: N45/13 and 12570-2021 approved by Stockholm Animal Ethical Board) to test the effect of *in vitro* αSyn fibrils by incubating on *ex vivo* hippocampal γ-oscillations. For brain extraction, mice were deeply anesthetized with isoflurane. The brain was dissected out and placed in ice-cold artificial cerebrospinal fluid (ACSF) modified for dissection containing (in mM): 80 NaCl, 24 NaHCO3, 25 glucose, 1.25 NaH2PO4, 1 ascorbic acid, 3 Na-pyruvate, 2.5 KCl, 4 MgCl2, 0.5 CaCl2, 75 sucrose and bubbled with carbogen (95% O_2_ and 5% CO_2_). Horizontal sections (350 μm thick) of the ventral hippocampi of both hemispheres were prepared with a Leica VT1200S vibratome (Leica Microsystems). Immediately after cutting, slices were transferred into a humidified interface holding chamber containing standard ACSF (in mM): 124 NaCl, 30 NaHCO3, 10 glucose, 1.25 NaH2PO4, 3.5 KCl, 1.5 MgCl2, 1.5 CaCl2, continuously supplied with humidified carbogen. The chamber was held at 37 °C during slicing and subsequently allowed to cool down to room temperature (∼22 °C) for a minimum of 1 hour. We selected four conditions to test the toxic effect of αSyn and BRICHOS R221E monomers on hippocampal γ-oscillations: Sham group (PBS, 0.1% Sodium Azide, pH 7.4), αSyn 500 nM, αSyn 1µM and αSyn 1µM + BRICHOS 1µM. All protein we produced in PBS buffer, 0.1% sodium azide, pH 7.4. We preincubated hippocampal slices in each condition for 30 minutes in a submerged incubation chamber containing ACSF. During the incubation, slices were supplied continuously with carbogen gas (5% CO_2_, 95% O_2_) bubbled into the ACSF. After incubation time the slices were transferred to the interface-style recording chamber for extracellular recordings (Chen et al., 2020; Kumar et al., 2023).

Recordings were performed with borosilicate glass microelectrodes filled with ACSF in hippocampal area CA3, pulled to a resistance of 3–6 MΩ. Local field potentials (LFP) were recorded at 32 °C in an interface-type chamber (perfusion rate 4.5 mL per minute). LFP gamma oscillations were elicited by kainic acid (100 nM). The oscillations were stabilized for 20 min before any recordings. Interface chamber LFP recordings were carried out by a 4-channel amplifier/signal conditioner M102 amplifier (Electronics lab, University of Cologne, Germany). The signals were sampled at 10 kHz, conditioned using a Hum Bug 50 Hz noise eliminator (Quest Scientific, North Vancouver, BC, Canada), software low-pass filtered at 1 kHz, digitized, and stored using a Digidata 1322 A and Clampex 10.4 programs (Molecular Devices, CA, USA). Power spectra density plots (from 60s long LFP recordings) were calculated in averaged Fourier-segments of 8192 points using Axograph X (Kagi, Berkeley, CA, USA). Gamma oscillations power was calculated by integrating the power spectral density between 20 and 80 Hz with the result representing average values taken over 1 min periods (Arroyo-García et al., 2021). One-way ANOVA test was used to compare the different conditions.

## Acknowledgement

The authors acknowledge financial support by FORMAS (A.A.), Swedish Society for Medical Research (A.A.), Alzheimer foundation (A.A.), Åke Wiberg foundation (A.A.), Magnus Bergvall foundation (A.A., H.B.), Åhlen foundation (A.A.), KI Research Foundation Grants (A.A.), Foundation for Geriatric Diseases KI (A.A.), Swedish Research Council (J.J.), Swedish Brain Foundation (J.J.) and JPco-fuND/EU PETABC 2020-02905/EC 643417 (H.B.). We thank Cecilia Mörman for commenting on the manuscript and Per Nilsson for providing mice.

## Author Contributions

A.A. and J.J. designed the study; L.A., R.K., L.E.A.-G., W.H.M., J.S.N., H.K., S.F., R.A., J.N., H.B. and A.A. performed experiments; L.A., R.K., L.E.A.-G., W.H.M., J.S.N., and A.A. analyzed data; L.A., L.E.A.-G., J.S.N. and A.A. visualized results; L.A. and A.A. wrote the original draft of the manuscript, all co-authors reviewed and edited the manuscript; A.A., J.J. and D.O. acquired funding and A.A., J.J. and D.O. supervised the study.

## Data availability statement

All data are available in the paper or as Supporting Information. Additional raw data can be obtained from the corresponding author upon request.

