## Supporting Information for "Specific inhibition of α-synuclein oligomer generation and toxicity by the chaperone domain Bri2 BRICHOS"

### Supporting Information Figures

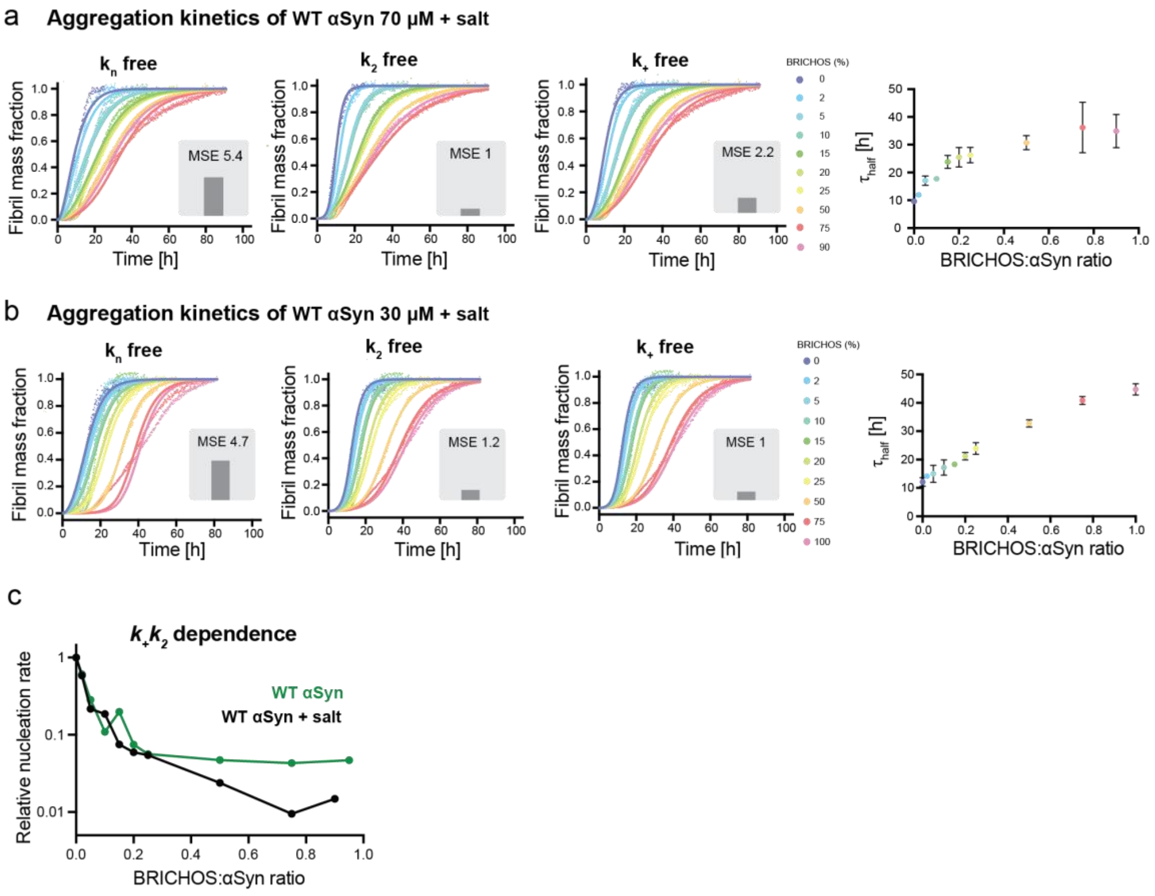


**SI Figure S1. BRICHOS inhibits αSyn aggregation in the presence of near-physiological salt concentration. (a,b)** Averaged and normalized aggreation kinetics of (a) 70 µM and (b) 30 µM αSyn (middle row) in the presence of 0 to 100 % BRICHOS monomers (color gradient from blue to red) in the presence of 154 mM NaCl. The aggregation traces were fitted to a secondary nucleation dominated model, where either $k_n$, $k_2$ or $k_+$ is the free fitting parameter. The MSE value corresponds to the normalized mean square error. The aggregation half-time increases with increasing BRICHOS molar ratios, at (a) 70 µM and (b) 30 µM αSyn concentration. **(c)** Dependence of $k_+k_2$ of the WT αSyn with and without salt on BRICHOS:αSyn ratio.

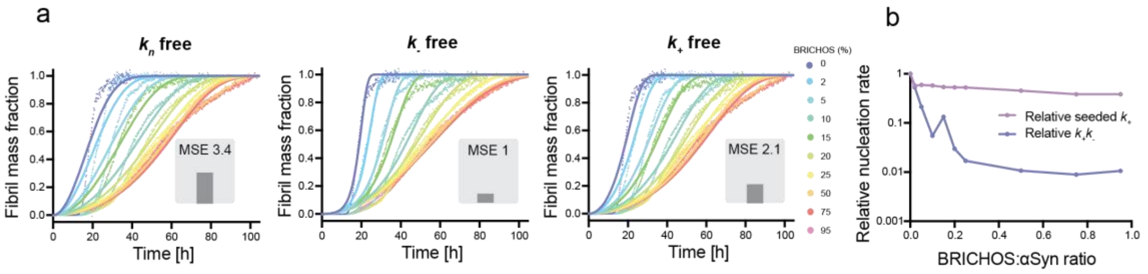


**SI Figure S2: αSyn aggregation kinetics fitted with a fragmentation-based model**. **(a)** Aggreation kinetics of 70 µM αSyn in the presence of 0 to 95 % Bri2 BRICHOS monomers (color gradient from blue to red) using the fragmentation dominated mechanism model. Individual fits (solid lines) of normalized and averaged aggregation traces (dots) of αSyn aggregation kinetics are shown in presence of BRICHOS where either $k_n$ , $k_-$ or $k_+$ is the free parameter. MSE values correspond to the normalized mean square errors. **(b)** Combined rate parameter $k_+k_-$ (where $k_+k_n$ is fixed as global constant) and $k_+$ extracted from highly seeded kinetics against relative BRICHOS concentration.

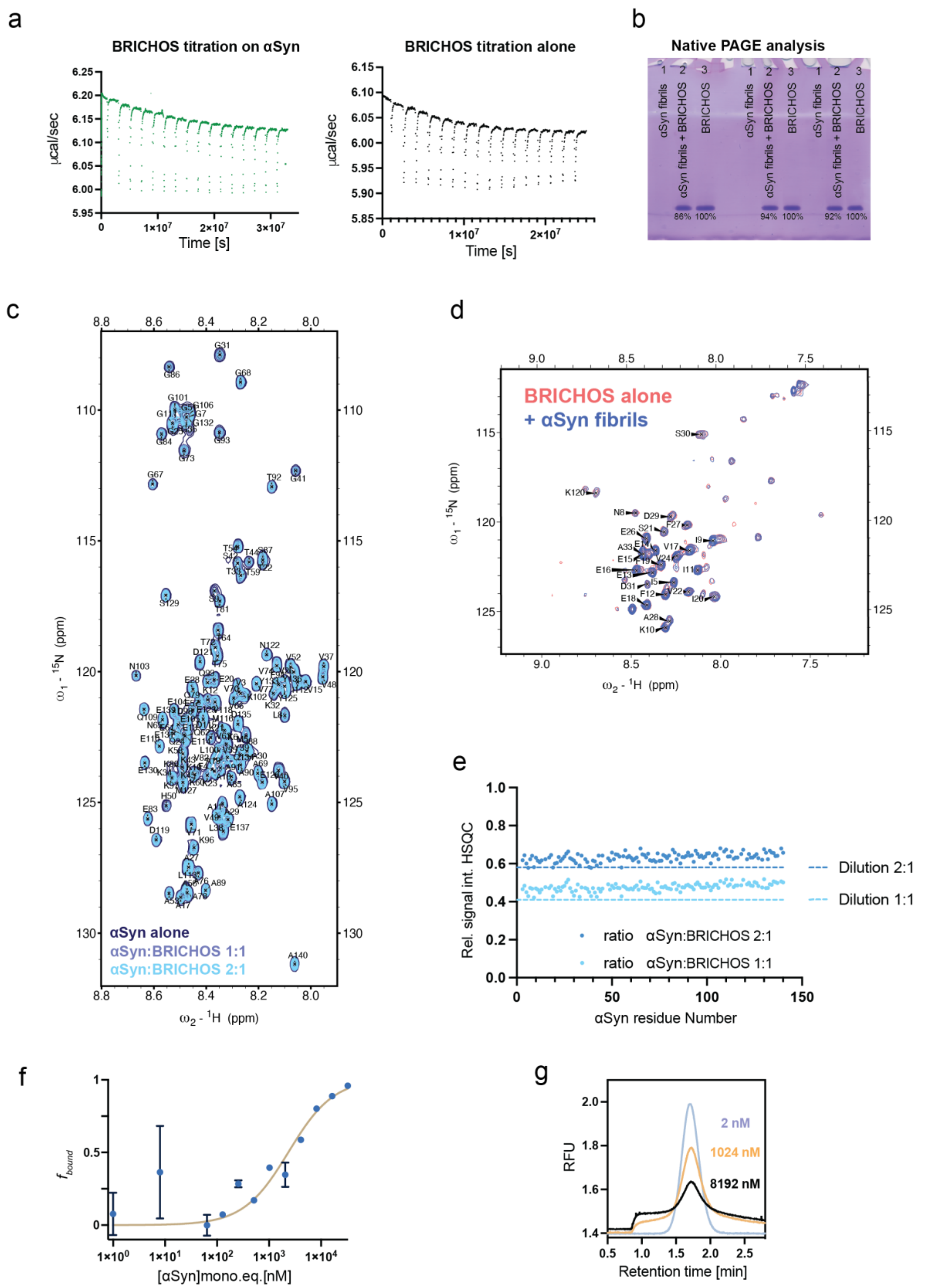


**SI Figure S3: BRICHOS binds to αSyn fibrils but not αSyn monomers**. **(a)** Isothermal titration calorimetry of BRICHOS monomers (syringe) on αSyn monomers (cell) (green) and control BRICHOS alone revealing the same heat dilution signal. **(b)** Native PAGE analysis of soluble BRICHOS using three replicates of BRICHOS incubated at equimolar concentration with αSyn fibrils. **(c,d)** 2D NMR $^{1}$H-$^{15}$N HSQC spectrum of (c) $^{15}$N-labeled αSyn monomers in presence of equimolar and submolar concentration of BRICHOS and (d) $^{15}$N-labeled BRICHOS in presence of equimolar concentration of sonicated αSyn fibrils. **(e)** Relative $^{1}$H-$^{15}$N HSQC signal intensities of $^{15}$N-labeled αSyn monomers in the presence of BRICHOS at different molar ratios. The dashed lines correspond to theoretical signal decrease after dilution, which agrees well with the observed signal attenuation. **(f)** Pre-mix FIDA titration of αSyn fibrils to BRICHOS-Alexa$^{488}$ represented by fraction of BRICHOS bound to the fibrils ($f_{bound}$) calculated from max- and minimal RFU values of the BRICHOS species. **(g)** Examples raw Taylograms of BRICHOS at 2, 1024, and 8192 nM of αSyn fibrils.
